# Origins of pandemic *Vibrio cholerae* from environmental gene pools

**DOI:** 10.1101/063115

**Authors:** B. Jesse Shapiro, Inès Levade, Gabriela Kovacikova, Ronald K. Taylor, Salvador Almagro-Moreno

## Abstract

Some microbes can transition from an environmental lifestyle to a pathogenic one^1–3^. This ecological switch typically occurs through the acquisition of horizontally acquired virulence genes^4,5^. However, the genomic features that must be present in a population prior to the acquisition of virulence genes and emergence of pathogenic clones remain unknown. We hypothesized that virulence adaptive polymorphisms (VAPs) circulate in environmental populations and are required for this transition. We developed a comparative genomic framework for identifying VAPs, using *Vibrio cholerae* as a model. We then characterized several environmental VAP alleles to show that, while some of them reduced the ability of clinical strains to colonize a mammalian host, other alleles conferred efficient host colonization. These results show that VAPs are present in environmental bacterial populations prior to the emergence of virulent clones. We propose a scenario in which VAPs circulate in the environment, they become selected and enriched under certain ecological conditions, and finally a genomic background containing several VAPs acquires virulence factors that allows for its emergence as a pathogenic clone.

## Main text

Numerous bacterial pathogens have emerged from environmental populations^1–3,6^. These virulent clones evolve through the acquisition of toxins and host colonization factors^4,5^. Given that the genes encoding these factors can often spread widely by horizontal gene transfer, it is surprising that only a limited number of pathogenic clones have emerged from any particular bacterial species. As a model of how environmental gene pools give rise to pandemic clones, we used *Vibrio cholerae*, a genetically diverse group of aquatic bacteria that include a confined phylogenetic group, the “pandemic genome” group (PG), that can cause the severe diarrheal disease cholera in humans^7,8^. Seven pandemics of cholera have been recorded to date, all caused by the PG group. The current pandemic is caused by strains of the El Tor biotype, and has spread across the globe in several waves of transmission^9^. Virulence in *V. cholerae* PG is mainly determined by two virulence factors: the cholera toxin (CT) and the toxin-coregulated pilus (TCP), which are encoded within horizontally acquired genetic elements, the CTXΦ phage and the Vibrio Pathogenicity Island-1 (VPI-1) respectively^10–12^. Both CTXФ and VPI-1 are always found in the PG group, however, they are also encoded in some environmental populations of *V. cholerae*^13–15^. Furthermore, even though some non-PG strains can cause gastrointestinal infections, only strains from the PG clade have ever emerged as a source of pandemic cholera^16^.

To investigate the evolutionary origins of pandemic clones of *V. cholerae* and the potential for their reemergence, we analyzed 43 *V. cholerae* genomes sequenced from clinical and environmental samples (Methods; Supplementary Table 1). These genomes span the known genetic diversity of *V. cholerae* (Supplementary Notes), and were divided into a primary dataset of 22 genomes and a replication dataset of 22 genomes, with one reference genome in common (Supplementary Table 2). In the primary dataset, we chose 7 PGs, including both classical strains, the source of the first six pandemics, and El Tor, to represent the genetic diversity of the pandemic group. We compared these with 15 non-clinical environmental genomes (EGs): 10 EGs from worldwide samples to include global diversity, and five sympatric isolates from the Great Bay Estuary (GBE) in New Hampshire, USA, a region with no recent history of cholera outbreaks^17^.

Consistent with the results of previous studies^7,9,17,18^, PGs form a distinct monophyletic group compared to EGs, based on the aligned core genome (Fig. 1a). Other than the PG group, there is little phylogenetic structure and the tree is star-like in both datasets (Supplementary Fig. 1). Reticulations in the phylogenetic network indicate recombination or homoplasies (repeated mutations in independent lineages at the same locus), consistent with a large, genetically diverse and recombining *V. cholerae* population^18,19^ (Supplementary Fig. 1). Given such a recombining population with mobile virulence factors, it remains puzzling why the ability to cause pandemic cholera is limited to the PG group.

**Figure 1.**
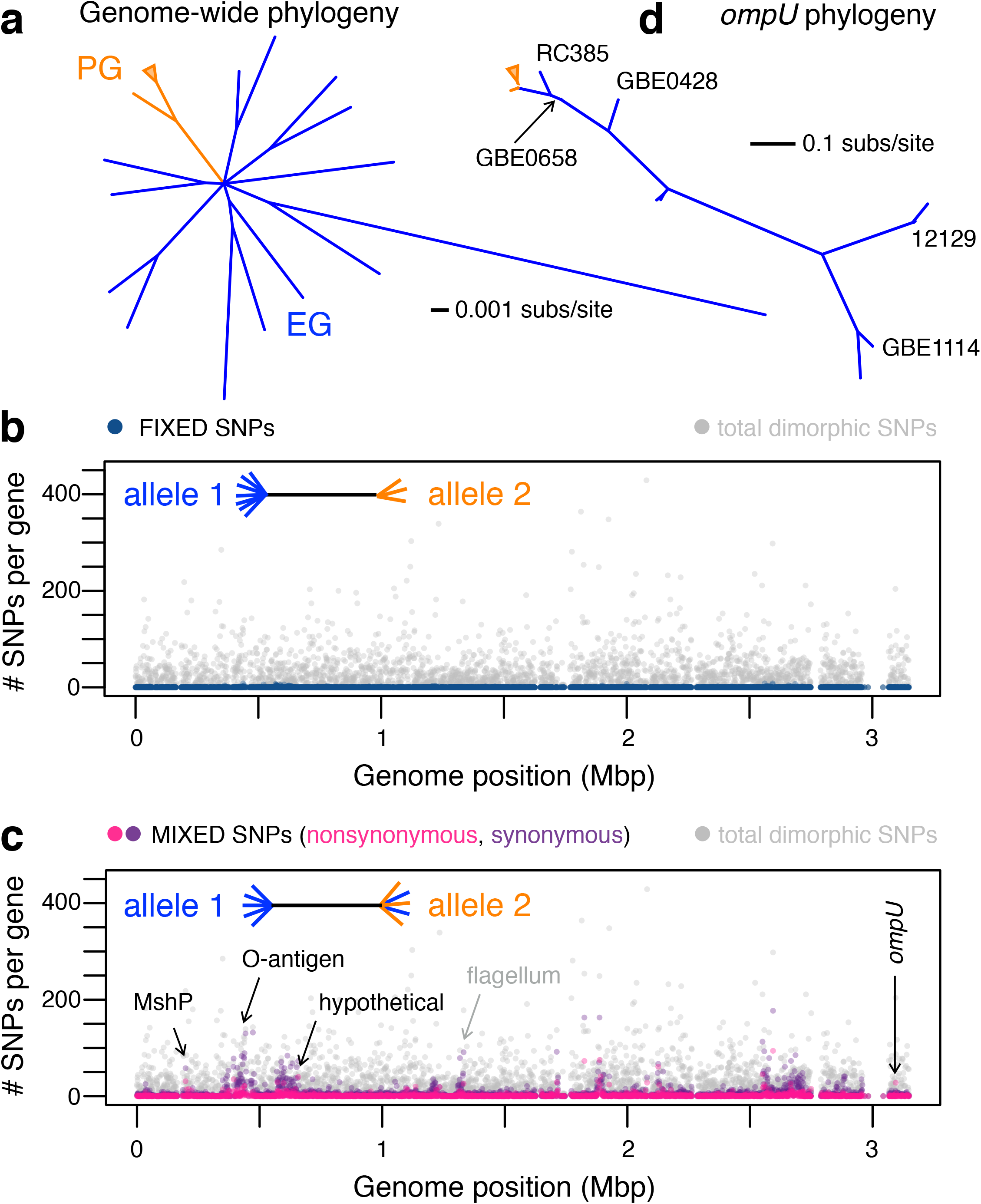
Comparative genomics reveals candidate virulence adaptive polymorphisms. **a**, Phylogeny of 22 *V. cholerae* genomes based on 1031 single-copy orthologs in the primary dataset. All branches have local support values >0.99 (based on FastTree's approximate likelihood ratio test) except for very short, deep internal branches (resulting in the star-like polytomy at the centre of the tree). Not all 22 genomes are visible because some have nearly identical sequences (*e.g*. 6 of the 7 PG genomes are nearly identical, shown as an orange triangle; GBE1173 and GBE1114 are nearly identical, as can be seen in Supplementary Fig. 1). **b**, Distribution of fixed SNPs across chromosome 1. (See Supplementary Fig. 3 for chromosome 2). Genome position is according to the MJ-1236 reference genome. SNP-free regions (*e.g*. near 3 Mbp, the locus of the integrative conjugative element) are part of the flexible genome, present in the reference but not the other 21 genomes. The schematic tree in the top left illustrates the fixed SNP pattern, in which one allele is present in PGs and a different allele in EGs. **c**, Distribution of mixed SNPs across the genome. The cartoon tree in the top left illustrates the mixed SNP pattern, in which one allele is fixed in PGs, and another allele is polymorphic among EGs, with some EGs containing the PG-like allele. Black arrows show candidate VAPs (Table 1). Grey arrow shows the flagellum as an example variable region not containing candidate VAPs. **d**, *ompU* phylogeny. All visible branches have local support values >0.9 except for the branch separating RC385 and GBE0658, the branch grouping MJ-1236 and O395 together, and the branch grouping HE09 and VL426 together.

To find unique features of PGs that could explain their pandemic success, we first searched for genes present in PGs but absent in EGs. Although there are a few genes and gene clusters which appear to be universally present in PGs, including CTXΦ and VPI-1, none of these are unique to PGs as they are also present in some EGs (Supplementary Table 3). This observation is consistent with previous work showing that these virulence genes are rapidly gained and lost in both EGs and PGs^7,18^. Therefore, there appear to be no gene families whose presence can easily explain the origin of pandemic cholera, nor are there strong boundaries to gene transfer between PGs and EGs.

Given the lack of PG-specific genes, we hypothesized that the environmental ancestor of the PGs had a particular genomic background containing alleles of core genes – which we term virulence adaptive polymorphisms (VAPs) – that served as “preadaptations” and enhanced its potential to give rise to pandemic disease. We started our search for VAPs by identifying SNPs in the aligned core of the 22 primary dataset genomes with one allele uniquely present in all PGs and a different allele uniquely present in all EGs. We called this a “fixed” SNP pattern. We identified 819 such fixed SNPs distributed across the genome (Fig. 1b; Supplementary Fig. 3). Some fixed SNPs could have contributed to the evolution of the pandemic phenotype in PG, while others could be selectively neutral hitchhikers on the PG genomic background. Using the McDonald-Kreitman test (Methods), we found evidence for genome-wide positive selection during the divergence of PGs from EGs, due to an excess of nonsynonymous changes at fixed sites, in both datasets (Supplementary Table 4). However, no individual gene showed evidence for selection after correcting for multiple tests (Methods), rendering it difficult to identify candidate VAPs using fixed SNPs. Nonetheless, fixed SNPs constitute only a modest fraction of possible SNP patterns (Supplementary Table 2), and VAPs could also exist at other SNP sites that might shed light in the evolutionary past of the ancestral PG.

We then defined and searched for a “mixed” SNP pattern, where PGs encode one fixed allele and EGs encode a “mix” of two alleles: one that is unique to EGs and also one that is fixed in PGs (Fig. 1c inset). We reasoned that the existence of PG-like alleles segregating in contemporary environmental populations of *V. cholerae* could be informative about: 1) pathogen emergence, because having an allele fixed in PG suggests that it could also have been present in the environmental ancestor of PG; and 2) the potential for pathogen reemergence, as PG-like alleles, and thus potential VAPs, are still circulating in the environmental gene pool. We identified 39,171 mixed SNPs in the primary dataset. Most genes contain few mixed SNPs (median of 3) but some genes contain dense clusters, resulting in a mean of 10.3 mixed SNPs per gene. The replication dataset contained greater genetic diversity, but showed the same pattern of a few genes containing many mixed SNPs (Supplementary Table 2). Clusters of genes containing many mixed SNPs are visible when plotted across the genome (Fig. 1c). Some of these clusters are known polymorphic regions of the genome and could be mutation hotspots containing SNPs not directly relevant to virulence adaptation. However, clusters of mixed SNPs do not visibly overlap with clusters of overall polymorphism (Fig. 1c), indicating that accumulation of mixed SNPs cannot be explained only by mutation hotspots.

To formally exclude mutation hotspots and focus on clusters of mixed SNPs shaped mainly by natural selection instead of mutation and drift, we considered only genes with an excess of nonsynonymous (NS) mixed SNPs relative to synonymous (S) mixed SNPs, which control for the baseline mutation rate. The mixed SNP pattern by definition groups PG-like EGs away from the other environmental strains and clusters them with the PGs. We reasoned that genes with an elevated NS:S ratio at mixed SNP sites were more likely to show phenotypic variations and have evolved under positive selection, possibly underlying preadaptations of the PG ancestor. Using a threshold of the number of mixed NS SNPs, NS:S, and *dN/dS* all two standard deviations above their genome-wide medians (Methods), we identified five genes as candidate VAPs in the primary dataset, three of which survived multiple hypothesis correction, and two of which (*ompU* and hypothetical gene VCD_001600) were also found in the replication dataset (Table 1). In contrast to the star-like genome-wide phylogeny (Fig. 1a), each of these five gene trees support one or more EGs grouping with PGs (Fig. 1d and Supplementary Fig. 4). Three additional VAPs – all hypothetical proteins – were identified in the replication dataset, suggesting the potential for other candidate VAPs to be identified with further sampling of genetically diverse environmental genomes (Supplementary Table 5).

**Table 1.**
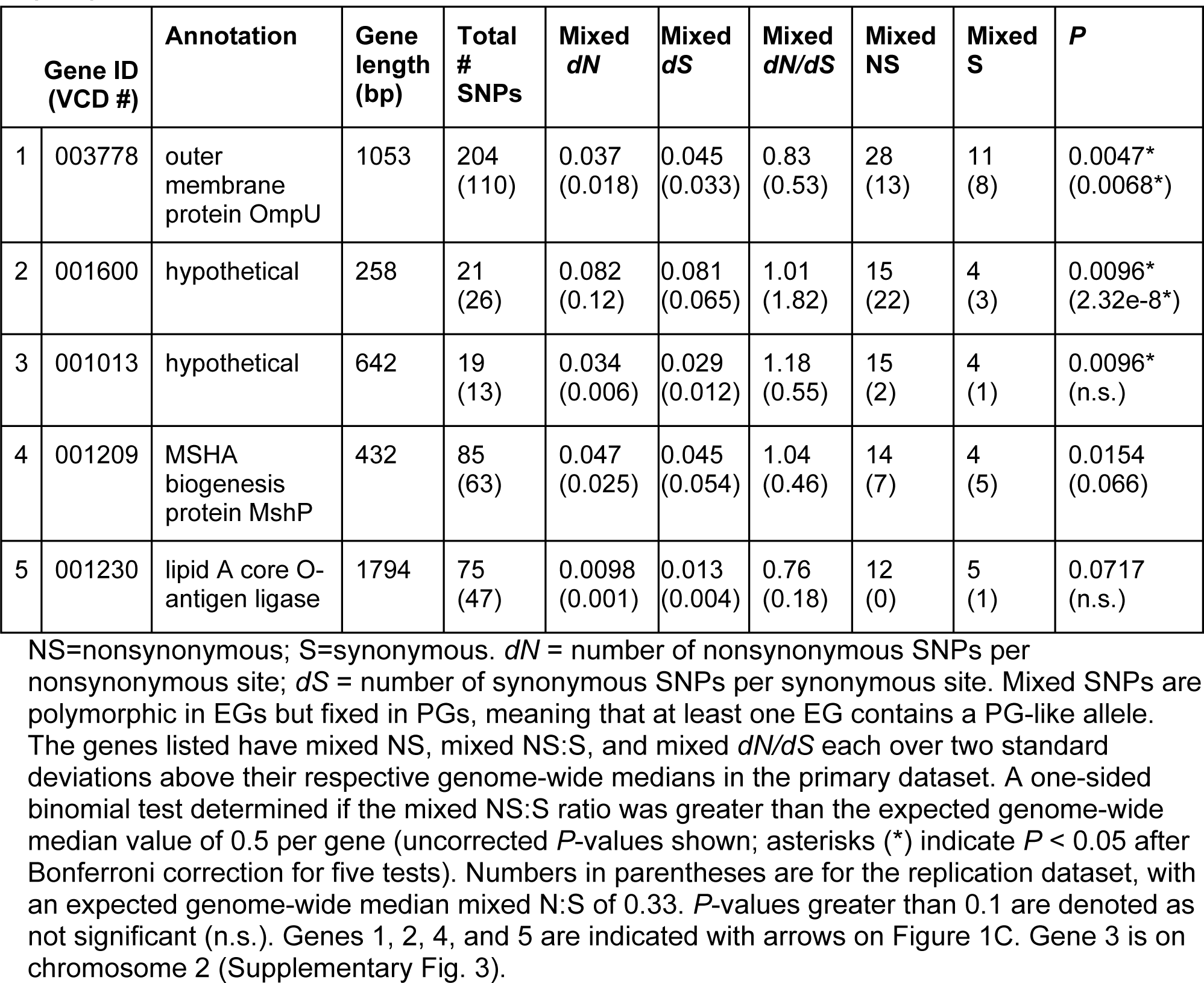
Characteristics of five predicted VAPs with an excess of nonsynonymous mixed SNPs.

Among the candidate VAPs, the gene with the most significant excess of mixed nonsynonymous SNPs in both datasets is *ompU* (Table 1). OmpU is an outer membrane porin that has been shown to play numerous roles in the intestinal colonization of *V. cholerae*, making it a compelling candidate for phenotypic characterization^20–23^. We observed that environmental strains RC385, GBE0658and GBE0428 have PG-like *ompU* alleles whereas most environmental strains, such as GBE1114, branch distantly from PG (Fig. 1d). We hypothesized that the PG-like alleles present in environmental strains might confer properties conducive to virulence. To test this, we constructed three different mutant strains each encoding one of three environmental alleles of *ompU* (all from sympatric GBE strains) into the background of N16961, a clinical strain from the current pandemic (Fig. 2). OmpU was detected on a protein gel stained with Coomassie blue and through immunoblot in all the constructed strains, indicating that all the strains effectively produce the environmental versions of the protein (Fig. 2a and Supplementary Fig. 6). The band size differences reflect the diversity in protein sizes among the three environmental OmpU variants (Supplementary Fig. 5 and Supplementary Table 6).

**Figure 2.**
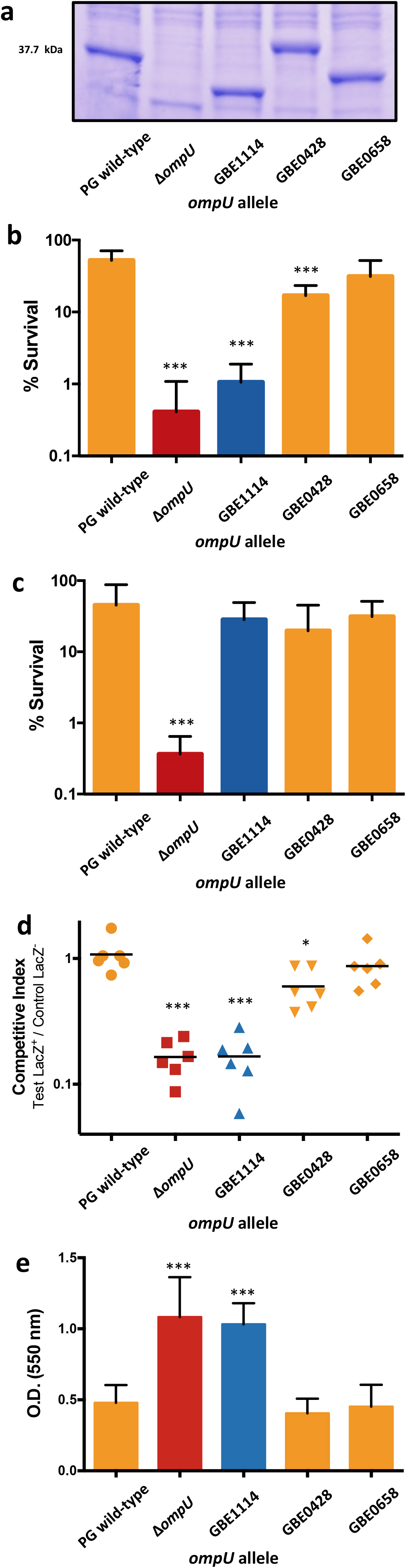
Phenotypic characterization of *ompU* alleles. **a**, OmpU production in clinical strains of *V. cholerae* encoding environmental alleles of *ompU*. Total protein lysates were run on a 16% Tris-glycine gel. OmpU bands were visualized after protein gels were stained with Coomassie blue. Three independent sets of protein lysates were examined and showed an identical band pattern. **b**, Survival of *ompU* mutants in the presence of bile (n=7) or **c**, polymyxin B (n=6). **d**, Colonization of the small intestine of *ompU* mutant strains (n=6). **e**, Biofilm formation of *ompU* mutant strains on an abiotic surface (n=15). Yellow bars and symbols, PG-like allele; red bars and squares, *∆ompU*; blue bars and triangles, EG-like allele. Center values represent the mean and error bars the standard deviation. Variance between the groups was similar. Statistical comparisons were made using Student's f-test. \**P* < 0.05, \*\*\**P* < 0.001

We performed three sets of experiments to compare the phenotypes conferred by EG and PG-like alleles of *ompU*. First, we determined the survival of these strains in the presence of 0.4% bile, as it has been previously shown that OmpU confers resistance to this antimicrobial compound^20^. The mutant strain encoding the *ompU* allele from the environmental strain GBE1114 (OmpU^GBE1114^) showed a similar survival rate in the presence of bile as a deletion mutant (Fig 2b). In contrast, OmpU^GBE0658^ show similar survival in the presence of bile as a PG strain (Fig. 2b). These experiments indicate that some environmental alleles of *ompU* confer properties beneficial for virulence. Second, we tested the survival of the mutants in the presence of polymyxin B, as OmpU also confers resistance against this antibiotic^21^. The ability to tolerate the antimicrobial effects of polymyxin B appears to be independent of which *ompU* allele is encoded by *V. cholerae*, as the three strains encoding environmental alleles of *ompU* had a similar survival rate (Fig. 2c). This experiment shows that OmpU^GBE1114^ is not simply a loss of function mutant, and is not equivalent to a knockout. Third, we determined the intestinal colonization of the *ompU* mutants by performing competition assays using the infant mouse model of human infection. We found that OmpU^GBE1114^ had a colonization defect similar to *∆ompU* whereas OmpU^GBE0658^ was able to colonize similarly to the strain with the wild-type PG allele (Fig. 2d). These results indicate that certain naturally occurring environmental alleles of *ompU* confer properties that provide an advantage to *V. cholerae* during or prior to host colonization, as would be expected for VAPs.

The *ompU^GBE1114^* allele appears to be maladaptive for intestinal colonization; however, its presence in several environmental isolates of *V. cholerae* prompted us to investigate its possible adaptive role in the environment (Fig. 2e). *V. cholerae* forms biofilms on the surface of biotic and abiotic environmental surfaces^24–26^, yet biofilm formation inside the host is thought to impair intestinal colonization^22,27^. Strains with deletions in *ompU* have been shown to form a more robust biofilm on abiotic surfaces^25^. We found that OmpU^GBE1114^ has higher biofilm formation than the strain encoding the wild-type PG allele, similar to the *ΔompU* strain (Fig. 2e). Both OmpU^GBE0658^ and OmpU^GBE0428^ formed biofilm similar to the strains with the wild-type PG allele (Fig. 2e). It therefore appears that there is an evolutionary trade-off between encoding the PG-like or EG-like alleles of VAPs, as they seem to confer mutually exclusive traits: either biofilm formation or bile resistance and host colonization. This suggests that environmental strains can be divided into subgroups which, due to their contrasting lifestyles, differ in their potential to give rise to pandemic clones.

We have determined that virulence adaptive polymorphisms are present in the environment, and shown how these VAPs can be identified, based on two independent sets of genomes. The top candidate VAP, *ompU*, was identified in both of our genomic datasets. Our experiments show that the *ompU* allele from some environmental strains, such as GBE0658, confers properties that allow for host colonization equally as efficient as alleles from clinical strains (Fig. 2). This leads to a natural question: Why have environmental strains with PG-like alleles not emerged as pandemic cholera strains? It appears that a variety of virulence adaptive genes and alleles are circulating in the environment, but only the PG group encodes the optimal combination of VAPs that allowed for pandemic potential (Fig. 3). We propose a conceptual model in which VAPs circulate in a diverse, recombining environmental gene pool, being maintained in the population through various biotic and abiotic selective pressures (Fig. 3a). A new ecological opportunity occurs, such as human consumption of brackish water or transient colonization of other animal hosts, which leads to the proliferation and gradual enrichment in the population of clones encoding a mosaic of VAPs (Fig. 3b). Finally, a genome encoding a critical combination of VAPs acquires key virulence factors allowing it to emerge as a virulent, potentially pandemic clone (Fig. 3c).

**Figure 3.**
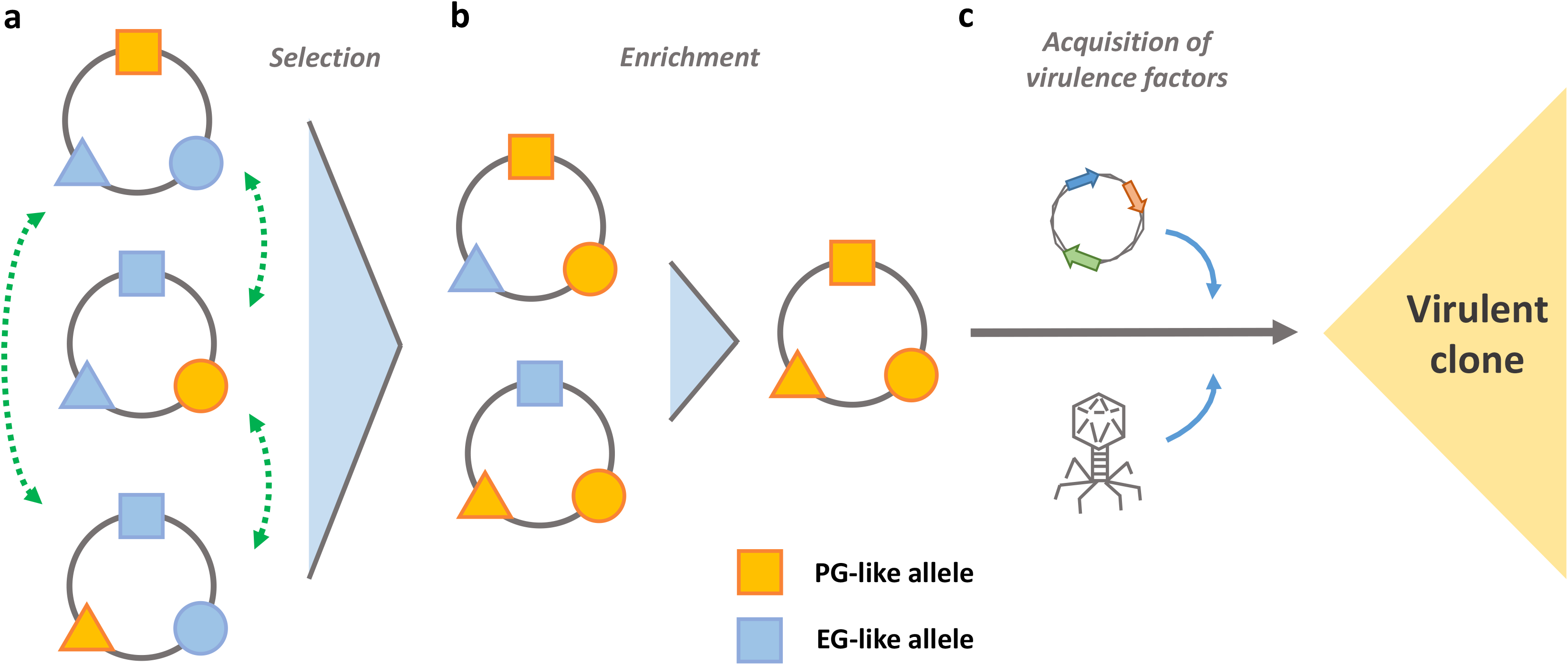
Model of pandemic clone emergence from an environmental gene pool. We propose a model that involves three events required for the emergence of pathogenic clones from environmental populations. **a**, selection of VAPs. Virulence adaptive alleles circulate in naturally occurring populations (orange symbols) and can be exchanged and mobilized through recombination (green dashed arrows). Ecological variation (*e.g*. temperature, nutrient availability, pH, etc.) leads to the selection of VAPs and an increase in their distribution in environmental populations. **b**, enrichment of clones. A new ecological opportunity occurs (human consumption of untreated waters, transient colonization of new environmental hosts, etc.) which leads to the proliferation and enrichment in the population of clones encoding a mosaic of VAPs. **c**, acquisition of virulence factors. A strain encoding a minimum set of VAPs required for host colonization acquires the virulence factors that are necessary to produce a successful infection and give rise to pandemic disease.

Our model posits that VAPs are circulating in the environment prior to the acquisition of key virulence factors. This is based on experimental evidence that a current pandemic strain cannot efficiently colonize the mammalian intestine without a PG-like *ompU* allele. If virulence adaptive alleles of *ompU* are indeed required prior to the acquisition of virulence factors, we would expect the same phenotypes of PG-like and EG-like *ompU* alleles in the genomic background of a more deeply branching PG isolate, such as classical *V. cholerae*. Indeed, we found that PG-like *ompU* alleles in the classical O395 background conferred efficient host colonization (Supplementary Fig. 7), which is consistent with an *ompU* VAP having played a role in the emergence of pandemic *V. cholerae*.

Our model further postulates that virulence adaptive alleles become enriched in the environmental population. Such enrichment would be made possible if these alleles provided a selective advantage in a newly available ecological niche, such as a human population consuming brackish water^28^. In previous work, we modeled an evolving, recombining microbial population that encounters a new ecological opportunity^29^. When adaptation to the new niche depends on few loci under positive selection in the niche, it is more likely for recombination to assemble the right combination of alleles in the same genome. These loci (akin to VAPs) could contribute additively or synergistically to fitness. For example, the PG-like *ompU* alleles confer a tenfold increase in fitness during host colonization (Fig. 2d). Other VAPs might contribute further to this enhancement in host colonization. As more loci are involved in adaptation, it becomes less likely to achieve the optimal combination. Assuming the *V. cholerae* population undergoes approximately 100 recombination events per locus per generation (6.5 recombination events for every point mutation)^19^, equivalent to a recombination rate of 10^−4^ in the modelled population of size 10^6^, an optimal combination of alleles at five loci could conceivably evolve, but seven loci is very unlikely^29^. Therefore, if virulence depended on five or fewer positively selected loci in the *V. cholerae* genome, the optimal combination of alleles would be expected to appear repeatedly in nature. Given suitable ecological opportunities, it is then plausible that pandemic *V. cholerae* could emerge multiple times, originating from outside the PG group. However, the number of loci that are sufficient for the emergence of a virulent strain remains unknown and if it was much greater than five, pathogen emergence would be naturally limited. We identified eight candidate VAPs in our two datasets, and if all eight are phenotypically confirmed to be VAPs, this number of VAPs (>5) might naturally limit pandemic clone emergence. Furthermore, these eight candidate VAPs passed stringent filters and we suspect there might exist additional VAPs in the genome, identifiable by further sampling and experimentation.

Here we have described a framework for identifying loci that are present in a natural population and confer properties beneficial for virulence prior to acquisition of essential virulence genes and host colonization. This framework could be applied to other bacterial pathogens that emerge as clonal offshoots from non-virulent relatives, including *Yersinia, Salmonella, Escherichia*, or other pathogenic *Vibrio* species^1–4,6^. Pathogens that emerged through clonal expansion limit our ability to dissect the genetic basis of their pathogenicity, because bacterial genome-wide association studies lack power when the phenotype of interest has evolved only once^30^. Our framework therefore provides a way forward to identify the genetic basis of virulence, even in pathogens that evolved through clonal expansion, and begin to assess the risk of pathogen emergence and reemergence from environmental gene pools.

## Methods

### Genome sequencing

DNA from clinical isolates (Bgd1, Bgd5, Bgd8, MQ1795^31,32^) and environmental isolates (GBE0428, GBE0658, GBE1068, GBE1114, GBE1173^17^) was extracted using the Gentra kit (QIAGEN) and purified using the MoBio PowerClean Pro DNA Clean-Up Kit. Multiplexed genomic libraries were constructed using the Illumina-compatible Nextera DNA Sample Prep kit following the manufacturer's instructions. Sequencing was performed with 250-bp paired-end (v2 kit) reads on the illumina MiSeq.

### Genome assembly

To exclude low-quality data, raw reads were filtered with Trimmomatic^33^. The 15 first bases of each reads were trimmed and reads containing at least one base with a quality score of <30 were removed. *De novo* assembly was performed on the resulting reads using Ray v2.3.1^34^.

### Genome alignment, annotation and SNP calling

We used mugsy v.1 r.2.2^35^ with default parameters to align the primary dataset of 22 *V. cholerae* genomes (Supplementary Table 1). From this alignment, we extracted dimorphic SNP sites and annotated genes according to MJ-1236 as a reference genome. We defined the core genome as locally colinear blocks (LCBs) with all 22 genomes present in the alignment. We replicated the alignment, annotation and SNP calling using 21 different *V. cholerae* genomes, mainly from Orata *et al*.^18^, plus the MJ-126 reference (Supplementary Table 1).

### Definition of orthologous groups

Genomes were annotated using the RAST web server (www.rast.nmpdr.org)^36^. Annotated genes were clustered into orthologous groups using OrthoMCL (http://www.orthomcl.org)^37^ with default parameters, yielding 2844 orthologous groups.

### Phylogenetic analysis

We constructed a core genome phylogeny using the concatenated alignment of 1031 single-copy orthologous protein-coding genes (present in exactly one copy in each of the 22 primary dataset genomes). Each protein sequence was aligned with Muscle^38^, and the concatenated alignment was used to infer an approximate maximum likelihood phylogeny with FastTree v. 2.1.8^39^ using default parameters (Fig. 1a). Individual gene trees (Fig. 1d) were built in the same way. We constructed a neighbour-net of the 22 genomes using SplitsTree v.4.10^40^, based on dimorphic SNPs from the mugsy genome alignment, excluding sites with gaps.

### Tests for selection

We conducted a genome-wide version of the McDonald-Kreitman test^41^ by first counting the number of fixed nonsynonymous (fn), fixed synonymous (fs), polymorphic nonsynonymous (pn), and polymorphic synonymous (ps) sites within each gene. We then summed these values across all genes (FN, FS, PN, and PS) and calculated the genome-wide Fixation Index, FI=(FN/FS)/(PN/PS). A fixation index greater than one suggests positive selection between the ingroup and outgroup (in this case, between EGs and PGs). However, care must be taken when computing a genome-wide FI because summing genes with different amounts of substitutions and polymorphism can result in FI>1 in the absence of selection^42^. We therefore performed 1000 permutations of the data, keeping the row totals (fn+fs and pn+ps) and column totals (fn+pn and fs+ps) constant and recomputing FI. We used the mean FI from the permutations as the expected value of FI under neutral evolution. To evaluate the hypothesis that the observed FI was higher than expected, suggesting positive selection, we computed a P-value as the fraction of permutations with FI greater or equal to the observed FI. We repeated the test using polymorphism from either the PG group or the EG group (Supplementary Table 3).

To identify individual genes under selection between PGs and EGs in the primary dataset, we restricted our search to 87 genes with fn > 0.68 1 and fn:fs > 1.48 (respectively two standard deviations about the genome-wide medians). We then calculated the gene-specific FI and assessed its significance with a Fisher exact test. We found no genes with FI significantly greater than one, after Bonferroni correction for 87 tests. Similarly, in the replication dataset, we restricted our search to 26 genes with fn > 1.5 and fn:fs > 1.70, none of which had significantly high FI after correction for multiple tests.

To identify genes with an excess of nonsynonymous mixed SNPs (likely due to selection for amino acid changes), we restricted our search to five genes with ≥ 12 NS mixed SNPs per gene and mixed NS:S > 1.78 (respectively two standard deviations above the genome-wide medians). We used a one-sided binomial test to assess whether the observed NS:S ratio for each gene was significantly greater than the genome-wide median NS:S ratio of 0.5 (after adding a pseudocount of one to both NS and S). Three out of the five genes had a significantly high mixed NS:S ratio (*P* < 0.05) after Bonferroni correction for five tests (Table 1). We repeated this procedure in the replication dataset, identifying genes with ≥ 18 NS mixed SNPs per gene and mixed NS:S > 1.65 (respectively two standard deviations above the genome-wide medians). We used a one-sided binomial test to assess whether the observed NS:S ratio for each gene was significantly greater than the genome-wide median NS:S ratio of 0.33 (after adding a pseudocount of one to both NS and S). The results of these tests are shown for genes also identified in the primary dataset (Table 1) and three additional genes identified in the replication dataset (Supplementary Table 5).

### Bacterial strains and plasmids

*V. cholerae* O395 and *V. cholerae* N16961 were used as wild-type strains of classical and El Tor biotypes respectively. Strains cultivated on solid medium were grown on LB agar; strains in liquid media were grown in aerated LB broth at 37°C. pKAS154 was used for allelic exchange^43^. When necessary, media was supplemented with antibiotics to select for certain plasmids or strains of *V. cholerae* at the following concentrations: gentamycin, 30µg/ml; kanamycin, 45µg/ml; polymyxin B, 50µg/ml; and streptomycin, 1 mg/ml.

### Strain construction

In-frame deletions and exchange of *ompU* alleles in both O395 and N16961 biotypes were constructed via homologous recombination^43^. For *ompU* deletions, PCR was used to amplify two 500 bp fragments flanking the *ompU* gene and to introduce restriction sites for cloning into the suicide vector pKAS154. For exchange of environmental *ompU* alleles, the respective allele was also amplified using the extracted DNA from each environmental strain. Restriction sites were introduced in the primers. The fragments were then cloned into a restriction-digested suicide plasmid, pKAS154, using a four-segment ligation for each environmental allele exchange mutants (plasmid, *ompU* flanking fragments and environmental *ompU* allele). The resulting plasmids were electroporated into *Escherichia coli* S17-1λpir. *E. coli* with the constructed inserts. Different sizes of *ompU* genes were confirmed, sequenced and compared to the genome sequences of environmental strains. For both deletion and exchange mutants, plasmids with the insert of interest were mated with wild-type *V. cholerae* O395 or N16961, and allelic exchange was carried out by selection on antibiotics as described previously^43^. For a more detailed description of allelic exchange please refer to Skorupski and Taylor 1998^43^. Potential mutants were screened using PCR: two primers flanking the deletion construct were used to amplify chromosomal DNA isolated from plated *V. cholerae*. The lengths of the PCR fragments were analyzed on 0.8% agarose gel for gene deletions and putative deletions and allele exchange were subsequently confirmed by DNA sequencing.

### OmpU visualization and immunobloting

Whole cell protein extracts were prepared from cultures grown for overnight at 37°C in a rotary shaker. The extracts were subjected to SDS-PAGE on 16% Tris Glycine gels (Invitrogen). OmpU bands were visualized after protein gels were stained by Coomassie blue overnight. Prior to staining the gels were transferred to nitrocellulose membranes using iBlot (Invitrogen). The membranes were blocked O/N in Tris-Buffered Saline, 3% BSA. Primary OmpU antibodies were diluted 1:10,000 in TBST (Tris-Buffered Saline, 0.5% Tween-20). Membranes were incubated with primary antibodies for 2 hours at room temperature. After incubation, the membranes were washed with TBST four times. Goat anti-rabbit secondary antibodies (BioRad) were diluted 1:10,000 in TBST and incubated for 30 minutes at room temperature. The membranes were washed 4 times with TBS (Tris-Buffered Saline). Reactive protein bands were detected via ECL (Amersham).

### Survival assays

*V. cholerae* strains were cultured overnight in LB broth at 37°C in a rotary shaker. Overnight cultures were diluted 1:100 in LB and grown to an OD600 of 0.5. Cells were pelleted and resuspended in either LB broth, LB containing 0.4% bile bovine (Sigma), or LB containing 1000U/ml of polymyxin B (Sigma). Cultures were incubated for 1h at 37°C in a rotary shaker. After incubation CFU/ml of each culture was calculated by plating dilutions in LB plates. Survival was calculated by comparing the number of CFU/ml in LB plus treatment versus LB. N≥6. No samples were excluded.

### Infant mouse competition assays

Overnight cultures were diluted 1:100. Each test strain was mixed in a 1:1 ratio with a *∆lacZ* reference strain. Four- to five-day-old CD-1 mice (*Mus musculus*) from several mixed litters were randomly inoculated orogastrically in blinded experiments with 50µl of the bacterial mixture. Sex of the animals was not inspected prior to inoculations. The intestines were harvested 24h post-inoculation and homogenized in 4ml of LB broth containing 10% glycerol. The mixtures were serially diluted and plated on LB agar plates supplemented with streptomycin and 5-bromo-4-chloro-3-indolyl-D-galactopyramoside (X-Gal)(40µg/ml). The competition indices were calculated as previously described by others, test (output CFUs/input CFUs)/reference (output CFUs/ input CFUs) and the sample size selected was appropriate for statistical analysis. No samples or animals were excluded. Animal work was approved by the Institutional Animal Care and Use Committee (IACUC).

### Biofilm assays

96-well plate assay. Cultures were incubated overnight at 30°C. 100µl of 1:100 dilutions of overnight cultures were placed per well in 96-well plates. Plates were left at 25°C for 24h. Liquid contents were discarded and plates were washed 2 times with LB. 200µl of 0.01% crystal violet was added per well and incubated at room temperature for 5 minutes. Liquid contents were discarded and plates were washed extensively with dH_2_O. After the plates were dry, biofilms were resuspended in 150µl of 50% acetic acid. Contents were transferred to a flat bottom dish and quantitated in a microtiter plate reader at OD550. Values were plotted using Prism software. N=15.

## Data availability

The nine genomes sequenced in this study (Bgd1, Bgd5, Bgd8, MQ1795, GBE0428, GBE0658, GBE1068, GBE1114, GBE1173) have been deposited under BioProject ID PRJNA349157 in NCBI GenBank under accession numbers SAMN05924900-SAMN05924908 (Supplementary Table 1). All other data supporting the findings of this study are available from the authors upon request.

## Acknowledgements

The authors would like to thank the anonymous reviewers for their thoughtful comments and suggestions. We would also like to thank Otto Cordero, Yves Terrat, Nicolas Tromas and Britney Privett for constructive comments on the manuscript. We thank Lawrence Shelven for his highly valuable technical assistance. BJS was supported by a Canada Research Chair and the Canadian Institutes for Health Research. RKT was supported by a National Institutes of Health grants AI039654 and AI025096. SAM was supported by startup funds from the Burnett School of Biomedical Sciences at the University of Central Florida and Dartmouth College's E. E. Just Postdoctoral Fellowship. This article is dedicated to the memory of Ronald K. Taylor.

## Author contributions

SAM conceived the study. BJS, RKT and SAM designed the study. IL sequenced genomes. BJS performed computational analysis. GK and SAM performed phenotypic characterization. BJS and SAM analyzed, interpreted data and wrote the article. All authors have read a version of the manuscript.

